# SARS-CoV-2 Nsp8 N-terminal domain dimerizes and harbors autonomously folded elements

**DOI:** 10.1101/2021.03.12.435186

**Authors:** Miguel Á. Treviño, David Pantoja-Uceda, Douglas V. Laurents, Miguel Mompeán

## Abstract

The SARS-CoV-2 Nsp8 protein is a critical component of the RNA replicase, as its N-terminal domain (NTD) anchors Nsp12, the RNA, and Nsp13. Whereas its C-terminal domain (CTD) structure is well resolved, there is an open debate regarding the conformation adopted by the NTD as it is predicted as disordered but found in a variety of complex-dependent conformations or missing from many other structures. Using NMR spectroscopy, we show that the SARS CoV-2 Nsp8 NTD features both well folded secondary structure and disordered segments. Our results suggest that while part of this domain corresponding to two long α-helices forms autonomously, the folding of other segments would require interaction with other replicase components. When isolated, the α-helix population progressively declines towards the C-termini, and dynamics measurements indicate that the Nsp8 NTD behaves as a dimer under our conditions.

## INTRODUCTION

The COVID-19 pandemic has currently (Mar. 12^th^, 2021) affected over 118 million persons and caused over 2.6 million deaths worldwide (https://coronavirus.jhu.edu/map.html). The virus responsible for the disease, SARS-CoV-2, is a coronavirus whose unusually long (30 kB) RNA genome is replicated by a rather sophisticated RNA polymerase. This polymerase complex is composed of several non-structural proteins (Nsp) including the RNA-dependent RNA polymerase, Nsp12, as well as Nsp7 and Nsp8, which embrace the RNA to promote progressivity^1^. Nsp8 serves as the platform onto which Nsp7 and Nsp12 bind, and cryo-EM studies have shown that it also contains binding regions to recruit the helicase Nsp13 within the replicase-transcription complex (RTC)^2,3^.

Due to this important role, a score of cryo-EM and crystallographic studies, detailed in **Sup. Table. 1**, have been applied to resolve the structure of Nsp8 in combination with Nsp7 (PDBs: 2AHM^1^, 6YHU^4^, 6M5I^5^, 6WQD^6^, 6XIP^7^) or Nsp7 plus Nsp12 (PDBs: 6M71^8^, 7BTF^8^, 6XQB^9^) and a model RNA template/primer (PDBs: 6YYT^2^, 7BW4^10^, 7B3B^11^, 7B3C^11^, 7B3D^11^, 7BV1^12^, 7BV2^12^, 7BZF^13^, 7C2K^13^) and Nsp13 (PDBs: 6XEZ^3^, 7CXM^14^, 7CXN^14^, 7CYQ^15^). In all these, the Nsp8 CTD appears well structured, adopting the same globular fold which interacts with Nsp7. Conversely, the NTD of Nsp8 shows a high degree of plasticity, with different conformations present in distinct contexts. Interestingly enough, bioinformatic analyses predict disorder in half of the Nsp8 NTD^16^, which may correlate with the absence of large portions of NTD residues in many crystallographic and cryo-EM structures (**Sup. Table 1**). This, together with the number of distinct non-structural proteins that anchor to Nsp8 to create the replicase, suggest that the Nsp8 behaves as an intrinsically disordered protein region (IDPR) that folds upon complexation. Indeed, an early crystallographic study of SARS Nsp8, whose NTD is identical to SARS CoV-2 Nsp8 except for the conservative Y15-->F substitution, showed that the NTD can adopt two strikingly different conformations in combination with a Nsp7 to form a hollow PCNA-like complex^1^. Building on the critical observation that the Nsp8 NTD integrates Nsp12, Nsp13 and RNA^12^, and its predicted intrinsic disorder and observed conformational plasticity prevent direct observation by crystallography or cryo-EM, we sought to characterize its conformation and dynamics using solution NMR spectroscopy. We advance that our results also provide a framework to rationalize the role of Nsp8 dimerization in the assembly of the replicase.

## RESULTS

### SARS CoV-2 Nsp8 NTD production and purification

Following purification by Ni^++^-NTA affinity, His tag cleavage and polishing with anion exchange chromatography (**Fig. S1B**), sample homogeneity was confirmed by SDS PAGE (**Fig. S1A**). The average yields were 5.0 mg/L from LB broth and 2.8 mg/L from minimal media for labeled samples.

### Nsp8 NTD solution secondary structure

The 2D ^1^H-^15^N HSQC spectrum of the Nsp8 NTD shows the excellent dispersion that is a hallmark of well-folded proteins for most signals (**Fig. 1A**). Closer inspection reveals a subset of narrower, more intense resonances corresponding to residues at the N- and C-termini. Spectral analysis led to the assignment of over 95% of the backbone ^13^Cα, ^13^CO, ^1^Hα, ^15^N and side chain ^13^Cβ chemical shifts; with ^1^H-^15^N assignments for all residues except Q56, R67, L59 and M62. The assigned chemical shifts have been deposited in the BMRB under accession code **50788**. The ^13^Cα conformational chemical shifts (Δδ) reveal two highly populated α-helical structures spanning residues 11-27 and 32-50 (**Fig. 1B**). Following residue 50, the helix continues but its population gradually declines, with the last stretch of residues (74-84) being chiefly disordered. The first ten residues are also disordered. The position and high population of the two α-helical segments was corroborated by ^1^Hα and ^13^CO conformational chemical shifts as well as ^1^HN-^1^Hα coupling constants (^3^J_HNHA_) (**Fig. S2**). This secondary structure, observed at 5 °C and quasi physiological pH, is maintained at 25 °C and 37 °C (**Fig. S3**). Regarding the residues linking the two α-helices, N28-G29-D30-S31, their ^13^Cα,^13^Cβ, ^13^CO, ^1^HN and ^1^Hα chemical shifts suggest that they adopt a type II’ tight turn ^17,18^.

**Figure 1:**
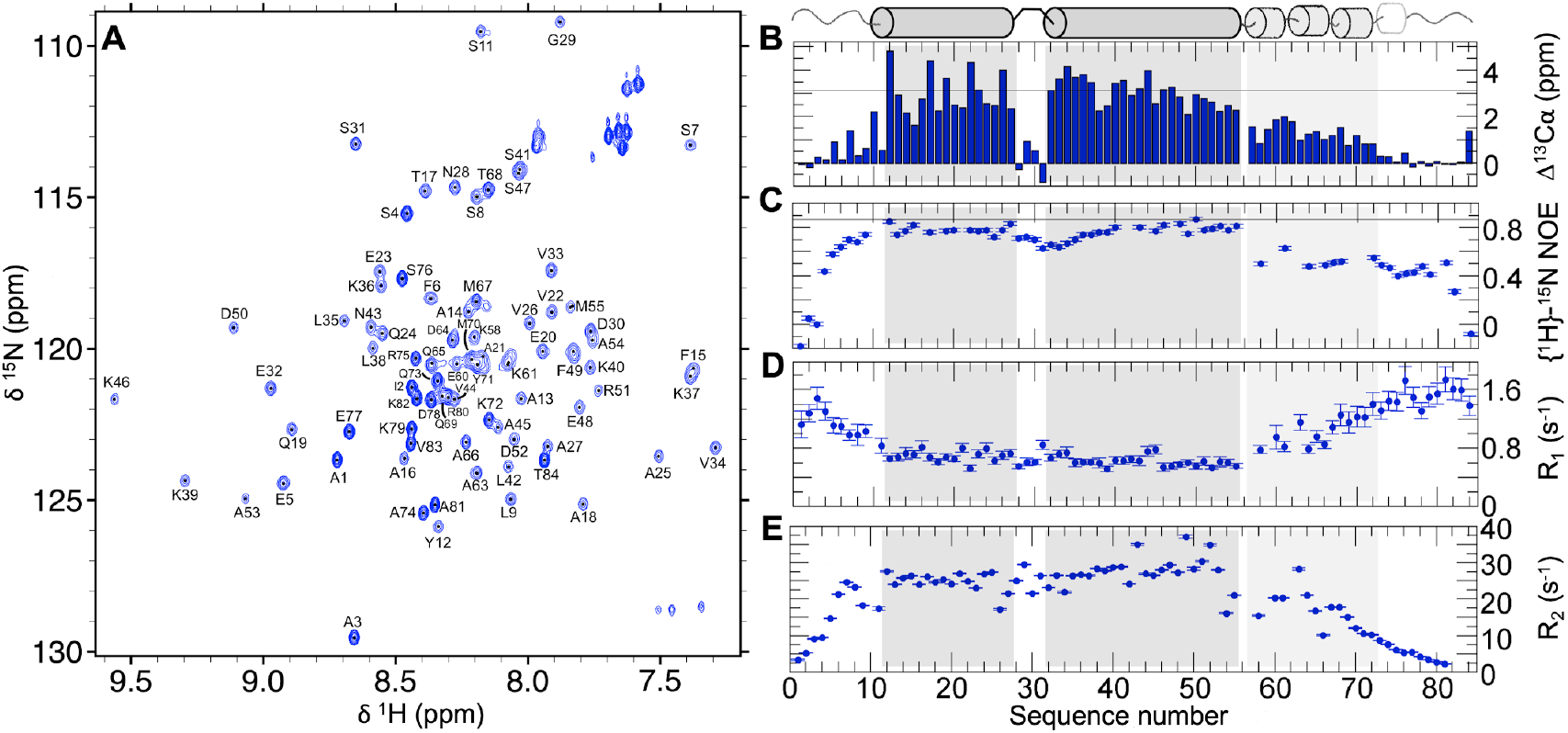
Isolated Nsp8 NTD adopts two central α-helices bordered by flexible termini. **A**. 2D ^1^H-^15^N HSQC spectrum of Nsp8 NTD recorded at 5.0 °C, in 50 mM NaCl, 10 mM KH_2_PO_4_, pH 6.1. Assigned backbone ^1^H-^15^N resonances are labeled. A set of sharper, more intense peaks with less ^1^H dispersion corresponding to N- and C-terminal residues are found amid a larger set of resonances with the wide range of ^1^HN chemical shifts characteristic of well folded proteins. **B**. ^13^Cα (blue bars) conformational chemical shifts (∆δ) reveal two highly populated *α*-helical structures spanning residues 11-27 and 32-50 (gray shading), as judged by the ∆δ ^13^C*α* value expected for 100% *α*-helix. Following residue 50, the helix continues, but its population declines in intensity (light gray shading). The thin line at 3.1 ppm marks the average Δδ^13^Cα value seen for fully populated α-helices. **C**. The heteronuclear {^1^H}-^15^N NOE. High ratios approaching the value of 0.86 (thin line) expected for full rigidity on ns/ps timescales are observed for the two main helical regions. **D**. Longitudinal (R_1_) and **E**. transverse (R_2_) relaxation rates gauge µs/ms timescale dynamics. Their relatively low (R_1_) and high (R_2_) values reflect stiffness and dimer formation.

### Solution Dynamics

The per-residue {^1^H}-^15^N NOE ratios, which are sensitive to dynamics on fast ps/ns timescales, are plotted in **Fig. 1C**. The residues composing the two α-helical segments show relatively high values averaging about 0.75. This indicates considerable stiffness, but is significantly below the theoretical ratio of 0.86 expected for complete rigidity, which is often observed in well-folded proteins. Ratios are lower in the first ten residues and decrease progressively beyond residue 60, reflecting increased mobility. The values for the residues 60 – 80 are in the neighborhood of 0.5, meaning that this segment is moderately mobile but significantly more rigid than a statistical coil which typically has values close to zero or even negative.

Residues 11 – 50 also show reduced longitudinal (R_1_) and elevated transverse (R_2_) relaxation rates which reflect dampened mobility on μs/ms timescales (**Fig. 1D**,**1E**). For this relatively rigid region, a correlation time (τ_c_) of 14.7 ns can be estimated which corresponds to an approximate molecular weight of 23 kDa in solution, assuming the protein behaves as an isotropic sphere^19,20^. Based on the findings of Bae *et al*. ^21^, a twenty residue disordered extension would add about 3 ns to the τ_c_, leading to a size estimate of 17.5 kDa. These values are roughly twice the 9.4 kDa MW per monomer based on the residue sequence x two monomers = 18.8 kDa. This indicates that Nsp8 NTD behaves as a dimer under these conditions.

## DISCUSSION

Nsp8 is an essential anchoring scaffold for the replication-transcription complex (RTC) of the SARS-CoV-2, whose NTD binds with the other critical components, particularly the RNA-dependent RNA polymerase Nsp12, the helicase Nsp13 and RNA. In these assemblies, the Nsp8 NTD adopts well-folded structures that contrast with bioinformatics analyses that predict this domain to be largely disordered^16^. In other many crystallographic or cryo-EM structures, large portions of the NTD are missing (**Table S1**) or adopt dramatically distinct conformations (**Fig. 2**), which evinces structural plasticity. In particular, the Nsp8 NTD of SARS-CoV, which is almost identical to SARS-CoV-2 Nsp8 NTD, adopts two strikingly different conformations when eight Nsp8 monomers combine with eight Nsp7 monomers to form a hexadecameric ring^1^ (**Figure 2A**). In one conformation, most of the Nsp8 NTD is absent whereas in the other, three α-helices are adopted (**Figure 2A**). By contrast, when combined with Nsp7, Nsp12 and RNA (with or without Nsp9 and Nsp13), the NTD of Nsp8 forms two α-helices which extend out like arms from the body of the RTC to embrace the RNA^2,3,8-15^ (**Figure 2B**). Two Nsp8 subunits are present per RTC, but they are not in contact. There is variation in the helix length, disorder and the position of the connecting turn among the reported structures. So what is the conformation of Nsp8’s NTD in isolation and what can it tell us about RTC assembly?

**Figure 2.**
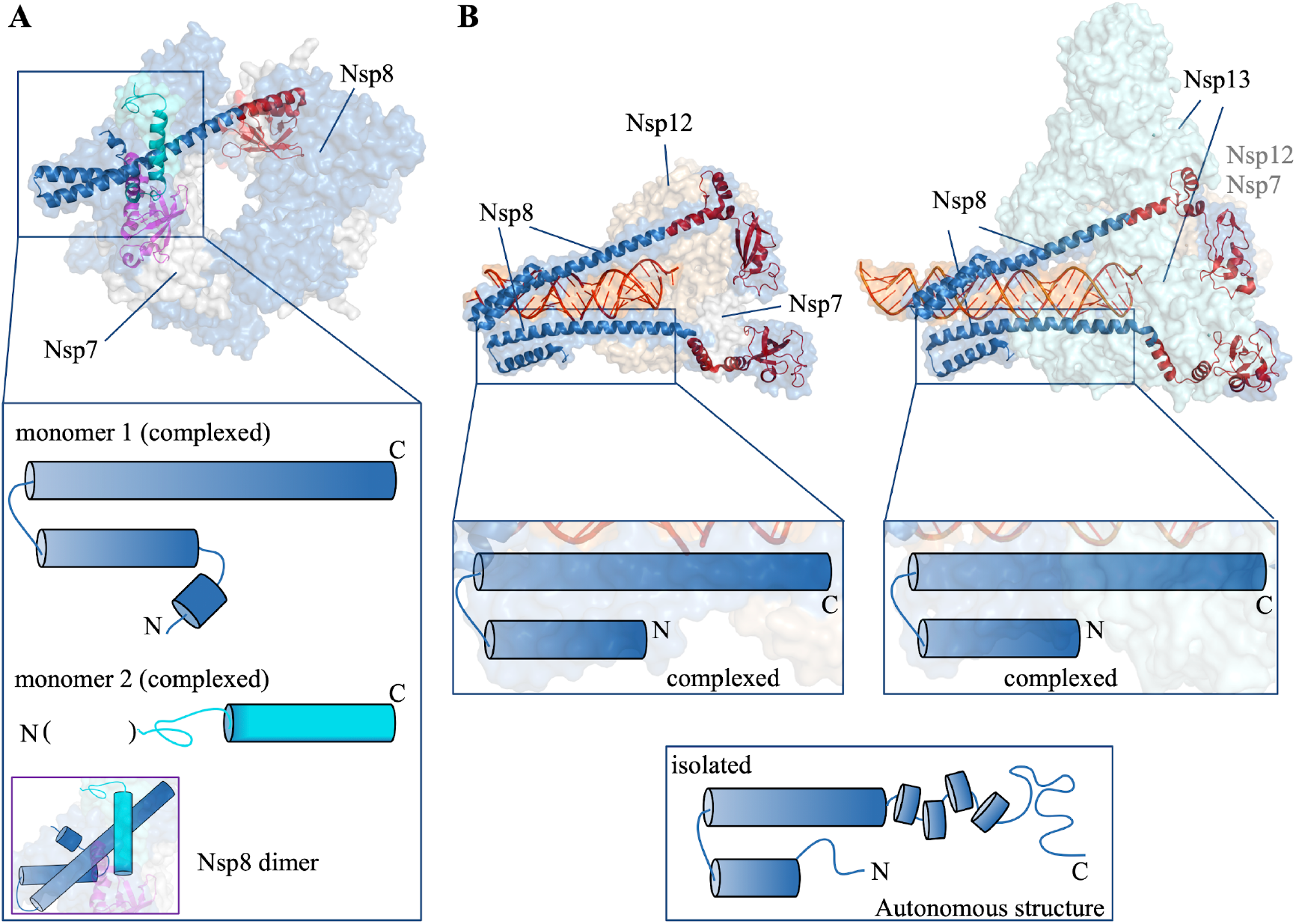
Conformational plasticity of the Nsp8 NTD. **A**. Two strikingly different conformations are adopted in Nsp8 NTD dimers in the hexadecameric ring adopted by combination of Nsp7 and Nsp8 dimers in SARS-CoV (PDB 2AHM)^1^. The zoomed box illustrates that one of the two monomers in the Nsp8 NTD dimers contains three helices, while in the second monomer part of the NTD is missing (see **Table S1**) but the remaining C-terminal part of the NTD preserves the helix. **B**. Two different structures of the RTC of SARS-CoV-2. The NTD of Nsp8 harbors interaction sites to bind RNA, Nsp12 (left, PDB 6YYT^2^ and Nsp13 (right, PDB 6XEZ)^3^, and adopts a helix-loop-helix conformation (zoomed regions). The bottom box illustrates an NMR-based 2D representation of the NTD in the absence of RNA, Nsp12 and Nsp13. Based on data shown in **Fig. 1**, a small yet rigid helix-loop-helix core forms autonomously, and is flanked by two N- and C-terminal segments. Whereas the N-terminal stretch is fully disordered, the C-terminal part possesses both nascent helical structure and a disordered segment. Nsp12 and Nsp13 bind to these C-terminal regions of the NTD, which become fully helical upon complexation.

One possibility is a concomitant structure formation upon binding event wherein an isolated, disordered Nsp8 NTD folds upon complexation. In line with this scenario, the Nsp8 forms distinct dimers with Nsp7 wherein the Nsp8 NTD folds in one type of dimer^1^ (**Fig. 2A**). In this structure and larger complexes, Nsp7 interacts with the CTD of Nsp8^2,3,8-15^ (**Fig. 2B**), which might suggest that the Nsp8 CTD then coaxes its NTD to fold. By contrast, here we show by NMR spectroscopy that the Nsp8 NTD, in the absence of other subunits and its CTD, exists as a folded dimer. This autonomous structure, consisting of two rather long, rigid α-helices spanning residues 11-50 and followed by a moderately-to-fully disordered segment (residues 51-84), might fold first and thus provide a foundation for anchoring Nsp7, Nsp12, Nsp13 and RNA to build up the RTC (**Fig. 2B**). Within this working model, the 51-84 residue segment could become fully helical as the complex grows (**Fig. 2B**).

One puzzling detail is the autonomous dimerization of the Nsp8 NTD, which could contribute to the formation of a hexadecameric ring structure^1^ (**Fig 2A**). However, the two Nsp8 NTD do not contact each other in recently reported assemblies of the RTC^2,3,8-15^ (**Fig 2B**). We can envision two possible scenarios. First, eight copies of Nsp8 and Nsp7 could really combine to form a hexameric ring, which would afford PCNA-like progressivity to the RTC as proposed by Zhai *et al*. ^1^ (**Fig 2A**). By contrast, if the physiological RTC were like the recent cryo-EM structures^2,3,8-15^ (**Fig. 2B**), then the dimerization of the Nsp8 NTD might initially serve to self-chaperone hydrophobic patches which evolve to form interactions with Nsp12, Nsp13 and RNA. The NMR data generated in this study represent valuable tools to test these scenarios by following changes in the assigned ^1^H-^15^N HSQC spectrum of Nsp8 NTD upon titration with other RTC components. As Nsp8 is the foundation of the replicase, these assignments shall also be key for identifying drug-like inhibitors for developing improved therapeutics by blocking Nsp8 NTD dimerization or associations with other replicase subunits. Beyond the replicase, these data can be used to map interactions with SARS-CoV-2 Orf6, a known partner^22^, which plays a key role in blocking the interferon response.

## MATERIALS AND METHODS

### Sample production and isotopic labeling

The gene coding for the Nsp8 NTD, whose sequence is G_0_A_1_IASE_5_FSSLP_10_SYAAF_15_ATAQE_20_AYEQA_25_VANGD_30_SEVVL_35_KKLKK_40_SLNV A_45_KSEFD_50_RDAAM_55_QRKLE_60_KMADQ_65_AMTQM_70_YKQAR_75_SEDKR_80_AKVT_84,_ was purchased from Genscript (New Jersey, NJ) with codons optimized for *E. coli* expression and subcloned in a pET28 derived vector containing in 5’ position an encoding sequence for thioredoxin followed by a hexa-histidine tag and a TEV-protease cleavage site. This construct was cloned in BL21star (DE3) cells. Afterwards, cells were grown in LB at 37 °C until OD_600_= 0.8, then the temperature was decreased to 25 °C, and 0.5 mM IPTG was added to induce the expression overnight. Uniformly isotopically labeled samples (^13^C + ^15^N) were obtained using a protocol derived from those described by Marley *et al*. ^23^ and Sivashanmugan *et al*. ^24^. Briefly, transformed cells were grown in 2 L LB media until OD_600_ = 2 - 3, then they were centrifuged, and the pellet resuspended in 0.5 L minimal medium containing ^13^C-glucose and ^15^NH_4_Cl. Cells were incubated at 37 °C for 1.5 h to enhance labeling. Next, the temperature was decreased to 25 °C before overnight induction with 0.5 mM IPTG.

The domain was purified by three chromatography steps. Briefly, after lysis by sonication, the obtained supernatant was purified on a HisTrap FF crude 5 mL column (Cytiva, Marlborough, MA) and eluted using an imidazole gradient whose initial and final concentrations were 10 and 500 mM, respectively. The eluted protein fusion was cleaved overnight at room temperature with TEV protease and dialyzed to eliminate imidazole. The cleaved sample was reloaded on the same column and the flowthrough was collected and applied to a Bio-Scale™ Mini Bio-Gel^®^ P-6 Desalting Cartridges (Bio-rad, Hercules, CA) for desalting. Finally, the sample was loaded on a 5 mL HiTrap Q HP anion exchange column (Cytiva, Marlborough, MA) at pH 8 and eluted, collecting the non-retained fraction. Homogeneity following purification was confirmed by gradient gel (4 - 20% acrylamide) SDS PAGE and NMR spectroscopy. Prior to NMR spectroscopy, the sample was transferred to a buffer containing 50 mM NaCl, 10 mM KH_2_PO_4_, pH 6.1. The spectra were referenced using DSS as the internal chemical shift standard. Spectra were recorded at 5.0 °C, except 2D ^1^H-^15^N HSQC and 3D HNCA spectra registered at 25 and 37 °C.

### NMR spectroscopy

A series of 2D ^1^H-^1^H TOCSY, ^1^H-^1^H NOESY (mixing time = 80 ms); ^1^H-^15^N HSQC, ^1^H-^13^C HSQC and 3D HNCO, HNCACO, HNCA, CBCAcoNH, CBCANH as well as 3D hNcocaNH and HncocaNH^25^ spectra were recorded on a Bruker Avance Neo 800 MHz (^1^H) spectrometer fitted with a cryoprobe and z-gradients. Spectral parameters are listed in **Sup. Table 2**. The spectra were transformed with Topspin 4.0.8 (Bruker Biospin) and were assigned by two independent operators using the aid of the program Sparky^26^. In addition, a series of 2D ^1^H-^15^N HSQC-based experiments were recorded to determine the {^1^H}-^15^N NOE, R_1_ and R_1_ρ relaxation rates in order to assess dynamics on ps/ns and μs/ms timescales, respectively. These spectra were recorded without non-uniform sampling and were transformed without applying linear prediction. For the {^1^H}-^15^N NOE ratios, peak integrals were measured using Topspin 4.0.8 and the uncertainties were calculated as the ratio of the noise (taken as the standard deviation of the integrals of several peakless spectral regions) times √2 to the peak integral measured without applying the NOE. R_1_ and R_1_ρ relaxation rates were obtained by using Bruker’s Dynamics Center program (version 2.6.3). For R_1_, uncertainties reported here are those obtained from the least squares fit. Transverse relaxation rates (R_2_) were calculated form R_1_ and R_1ρ_ and the uncertainties in R_2_ were obtained by error propagation.

## ACKNOWLEDGEMENTS

MM is a Ramón y Cajal Fellow of the Spanish AEI-Ministry of Science and Innovation (RYC2019-026574-I), and a “La Caixa” Foundation (ID 100010434) Junior Leader Fellow (LCR/BQ/PR19/11700003). Funded by project COV20/00764 from the Carlos III Institute of Health and the Spanish Ministry of Science and Innovation to MM and DVL. NMR experiments were performed in the “Manuel Rico” NMR Laboratory (LMR) of the Spanish National Research Council (CSIC), a node of the Spanish Large-Scale National Facility for Biomolecular NMR (ICTS R-LRB).

---

**Supporting Figure 1.**
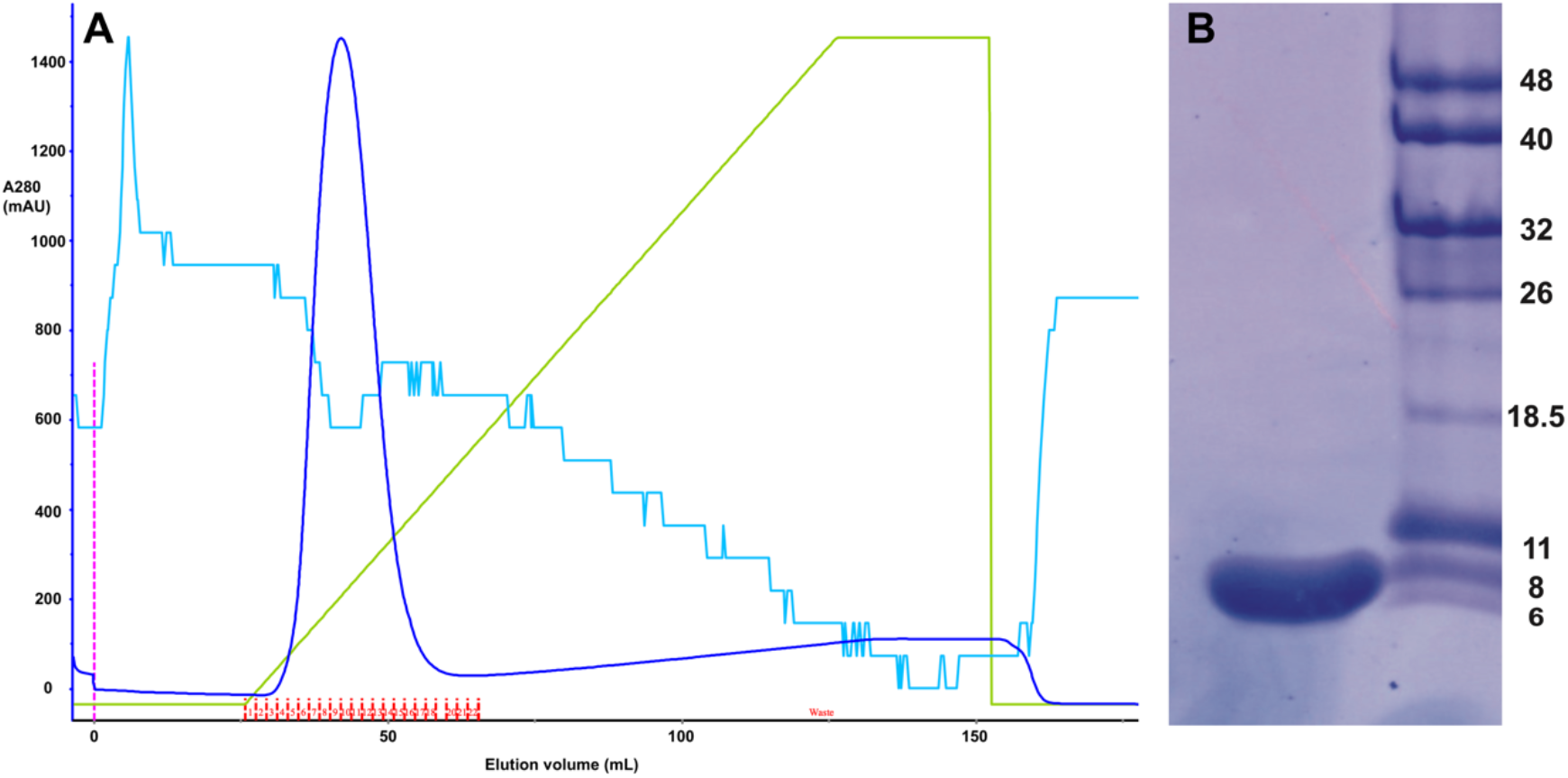
Purification of Nsp8 NTD. **A**. Chromatogram of the elution of ^13^C-^15^N Thioredoxin-His_6_-TEVcleavage sequence-Nsp8NTD from the Ni^++^ NTA column. The blue trace corresponds to absorbance at 280 nm, the green trace marks the imidazole gradient, which ranged from 10 mM to 500 mM, the red lines mark the collected fractions, the magenta line indicates the injection point and the light blue line represents the conductivity. **B**. Gradient (4 – 20% arcylamide) PAGE-SDS gel of the final purified ^13^C,^15^N Nsp8 NTD. The numbers on the right indicate the size, in kDa, of the molecular weight markers.

**Supporting Figure 2:**
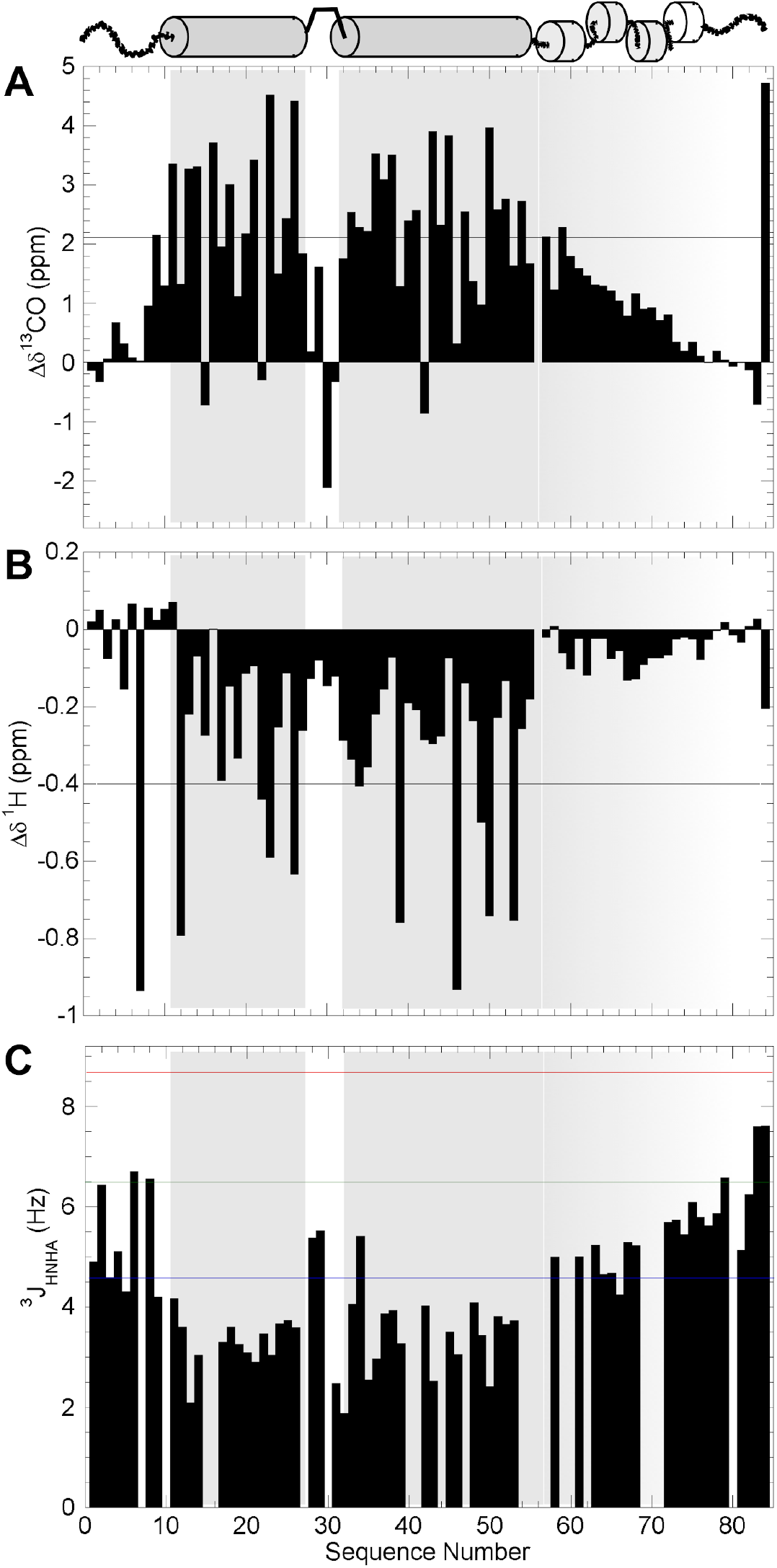
NMR corroboration of the presence of two α-helices linked by a tight turn and flanked by disordered N- and C-termini. Conformational chemical shift values, Δδ^13^CO (panel **A**), and Δδ^1^H (panel **B**), and ^3^J_HNHA_ (panel **C**) coupling constant values are plotted versus sequence number. The lines mark the values expected for α-helix in panel **A** and **B**; in panel **C**, the red, green and lines represent the values expected for β-sheet, statistical coil and α-helix, respectively.

**Supporting Figure 3.**
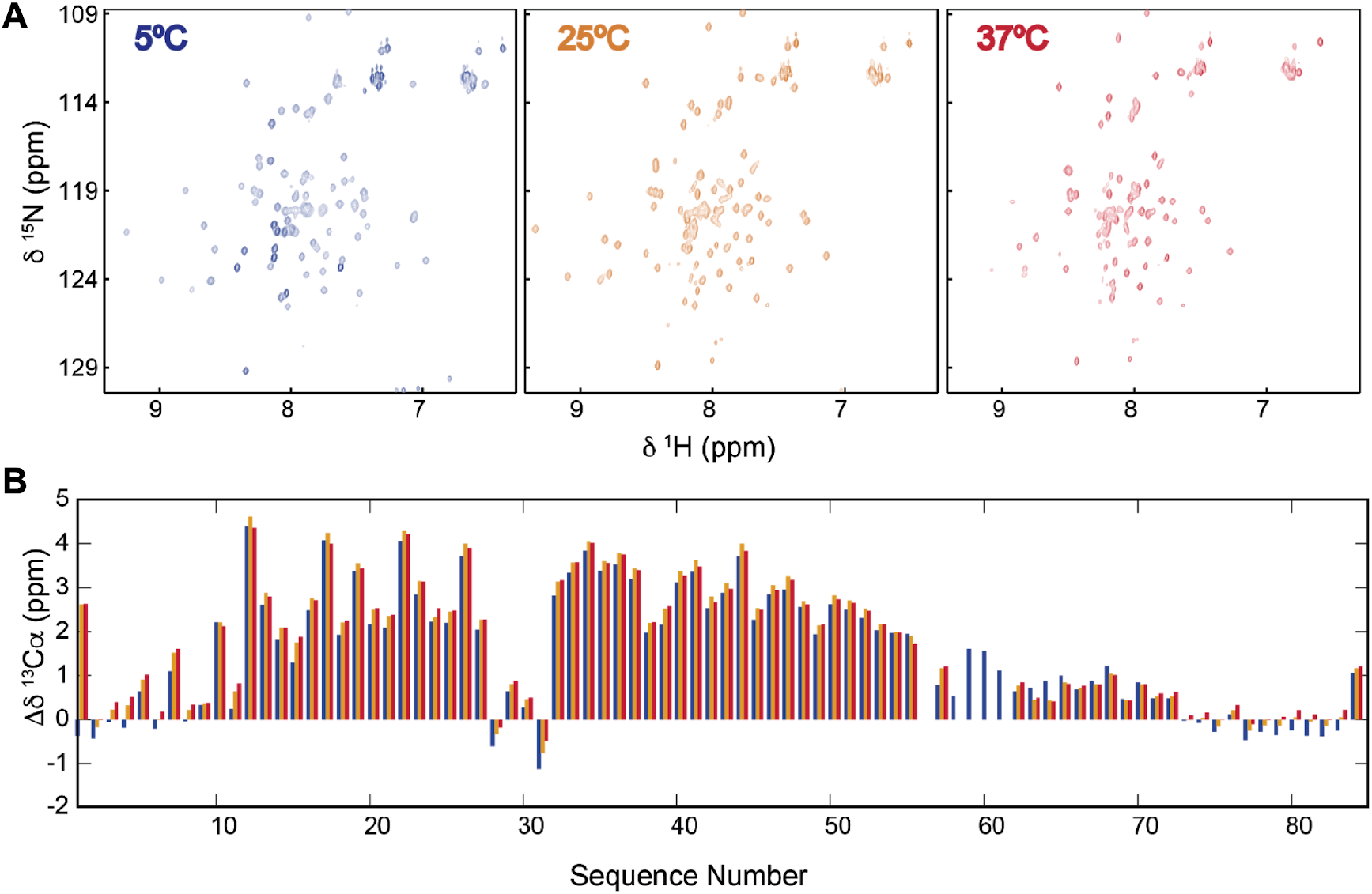
The Nsp8 NTD preserves its autonomous structure at physiological temperature. **A**. 2D ^1^H-^15^N HSQC spectra of ^13^C,^15^N-labeled Nsp8 NTD at 5.0 °C (left panel) 25.0 °C (middle) and 37.0 °C (right panel). Although some peaks lower in intensity due putatively to exchange with ^1^H_2_O, native signals are generally preserved at 37 °C. **B**. ^13^Cα conformational chemical shifts measured on the basis of 3D HNCA spectra at 5.0 °C (blue bars), 25.0 °C (gold bars) and 37.0 °C (red bars). The very small changes indicates that the content of *α*-helix scarcely changes over this temperature range.

**Supporting Table 1:**
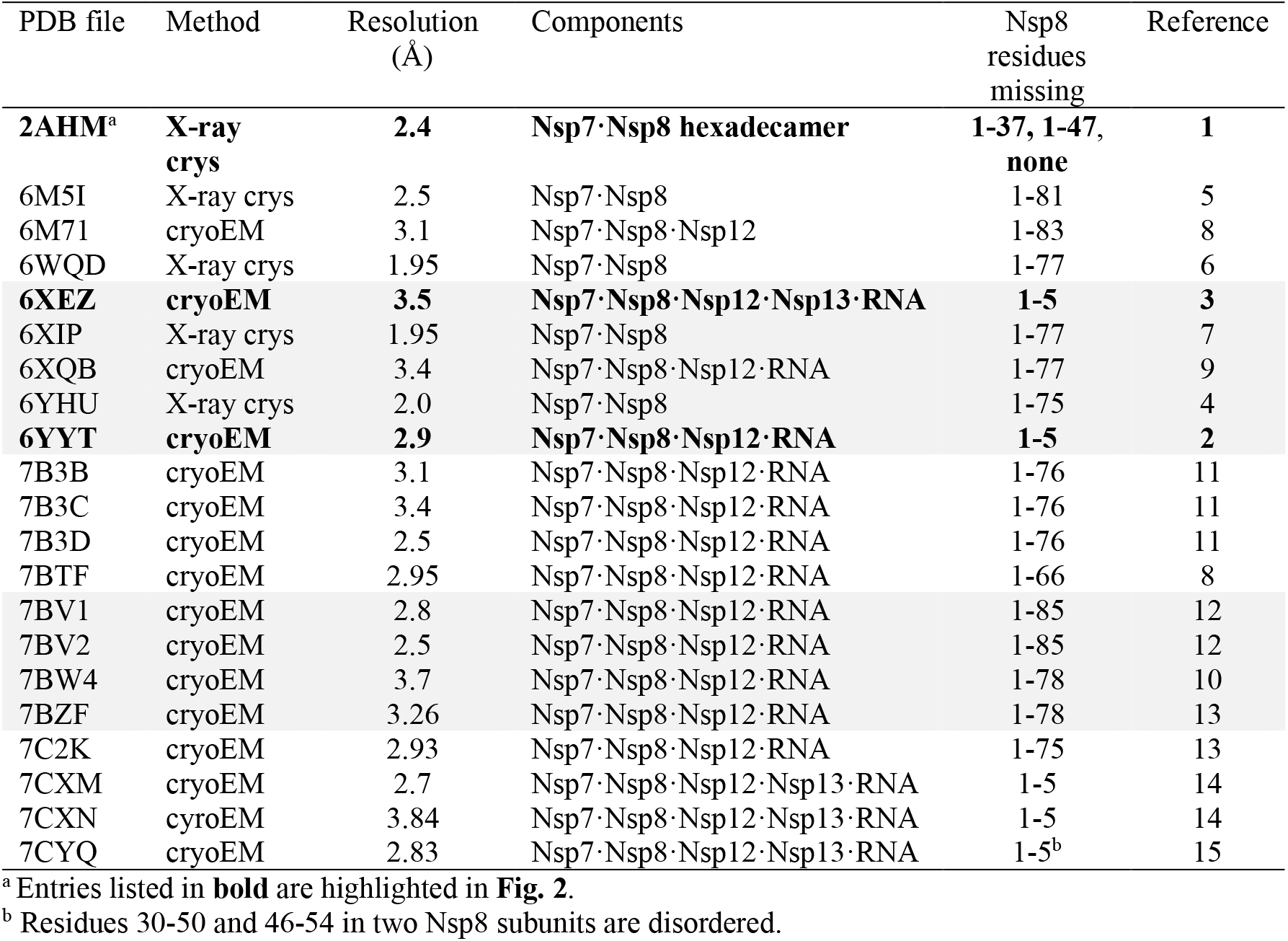
Previous medium/high resolution structural studies of Nsp8

**Supporting Table 2:** NMR Spectral Parameters

| Experiment | Number of Scans | Sweep Width (ppm) | Matrix |
| --- | --- | --- | --- |
| **Assignment** |  |  |  |
| 1D $^1\text{H}$ | 64 | 12 $^1\text{H}$ | 32k |
| 2D $^1\text{H}$-$^{15}\text{N}$ HSQC | 4 | 11 $^1\text{H}$ x 20 $^{15}\text{N}$ | 2k x 256 |
| 2D $^1\text{H}$-$^{13}\text{C}$ HSQC | 4 | 11 $^1\text{H}$ x 80 $^{13}\text{C}$ | 1k x 512 |
| 3D HNCO<sup>a</sup> | 8 | 10 $^1\text{H}$ x 22 $^{15}\text{N}$ x 12 $^{13}\text{CO}$ | 2k x 64 x 64 |
| 3D HNCA<sup>b</sup> | 8 | 10 $^1\text{H}$ x 22 $^{15}\text{N}$ x 25 $^{13}\text{C}\alpha$ | 2k x 64 x 128 |
| 3D CBCAcoNH<sup>c</sup> | 8 | 10 $^1\text{H}$ x 23 $^{15}\text{N}$ x 55 $^{13}\text{C}$ | 2k x 64 x 128 |
| 3D HNCACB | 16 | 10 $^1\text{H}$ x 23 $^{15}\text{N}$ x 80 $^{13}\text{C}$ | 2k x 64 x 104 |
| 3D HNHA | 16 | 10 $^1\text{H}$ x 10 $^1\text{H}$ x 23 $^{15}\text{N}$ | 2k x 64 x 64 |
| 3D HNcaCO | 8 | 10 $^1\text{H}$ x 23 $^{15}\text{N}$ x 16 $^{13}\text{CO}$ | 2k x 64 x 88 |
| **Relaxation** |  |  |  |
| 2D $T_1$<sup>d</sup> | 8 | 10 $^1\text{H}$ x 23 $^{15}\text{N}$ | 2k x 256 |
| 2D $T_{1\rho}$<sup>e</sup> | 8 | 10 $^1\text{H}$ x 23 $^{15}\text{N}$ | 2k x 256 |
| 2D $\{^1\text{H}\}$-$^{15}\text{N}$ NOE<sup>f</sup> | 16 | 10 $^1\text{H}$ x 20 $^{15}\text{N}$ | 2k x 400 |

<sup>a</sup> Non-uniform sampling (NUS) applied with 30% sampling.
<sup>b</sup> NUS applied with 35% sampling.
<sup>c</sup> NUS applied with 50% sampling.
<sup>d</sup> Eight spectra recorded with delays of 8, 1600, 20, 800, 100, 460, 60 and 240 ms.
<sup>e</sup> Eight spectra recorded with delays of 8, 600, 36, 300, 76, 200, 100 and 156 ms.
<sup>f</sup> Recycle delay = 10 seconds.

## Notes

https://bmrb.io

### Competing Interest Statement

The authors have declared no competing interest.

